# Multiple stressors in river networks: local and downstream effects on freshwater macroinvertebrates

**DOI:** 10.1101/2024.09.27.615445

**Authors:** Gemma Burgazzi, Noël P.D. Juvigny-Khenafou, Verena C. Schreiner, Alessandro Manfrin, Jonathan Jupke, Jeremy Piggott, Eric Harvey, Akira Terui, Florian Leese, Ralf B. Schäfer

**Affiliations:** iES Landau, Institute for Environmental Sciences, RPTU Kaiserslautern-Landau, Landau in der Pfalz, Germany; ALPSTREAM Group, Department of Life Sciences and Systems Biology, University of Turin, Turin, Italy; Institute of Aquaculture, University of Stirling, Stirling, Scotland (UK); Faculty of Biology, University of Duisburg-Essen, Essen, Germany; Research Center One Health Ruhr, University Alliance Ruhr, Essen, Germany; Zoology Department, School of Natural Sciences, Trinity College Dublin, Dublin, Ireland; Dépt des Sciences de l’environnement, Université du Québec à Trois-Rivières, Canada; Department of Biology, University of North Carolina at Greensboro, Greensboro, NC, USA

**Keywords:** flow intermittency, light pollution, intermittent rivers, artificial light at night, mesocosm, benthos, drift, meta-ecosystem

## Abstract

River networks are complex ecosystems characterized by a continuous exchange of material and energy through longitudinal gradients. These ecosystems are threatened by various human-induced stressors, which frequently co-occur and may interact in complex ways, potentially triggering cascading effects in the river network. Aiming at assessing single and combined effects of flow intermittency and light pollution on macroinvertebrate communities, we performed a multiple stressors experiment in 18 flow-through mesocosms. Each mesocosm was designed to mimic a simplified river network, with two upstream tributaries merging downstream, to assess both local and cascading effects. The experiment was performed in Summer 2021 for seven weeks (26 days of colonization, 23 days of treatment), applying the stressors either separately or combined in the upstream sections, in a randomized block design. Flow intermittency was simulated as the ponded phase of the drying process, whereas light pollution was applied with LED strips (set at 10 lux) that automatically turned on at sunset and off at sunrise. Drifting macroinvertebrates were sampled weekly during the treatment phase, and benthic macroinvertebrates at the end of the treatment phase. Both stressors individually applied had negative effects on the benthos, whereas drift decreased with flow intermittency and increased with light pollution. When combined upstream, stressors showed dominant effects of flow intermittency on the benthos and interactive effects on the drift. The effects of the single stressors and their interactions propagated along the river network, with stronger downstream effects when stressors co-occurred upstream. These findings showed that the spatial distribution of multiple stressors along the river network can affect their resultant downstream effects, highlighting the importance of framing multiple stressors research in a spatial context. Considering the pressing needs of the growing human population, our results represent a step forward in anticipating cumulative stressors effects, informing efficient conservation strategies for protecting freshwater ecosystems.

## 1. Introduction

A growing number of anthropogenic stressors are affecting both terrestrial (Saatchi et al., 2021; Smith et al., 2016) and aquatic ecosystems (Ormerod et al., 2010; Reid et al., 2019), posing new challenges to biodiversity conservation. Among others, freshwater ecosystems are the most severely impacted by human presence (Dudgeon, 2019; Tickner et al., 2020). In Europe alone, approximately 60% of rivers are affected by at least one stressor (Schinegger et al., 2012) and the co-occurrence of multiple stressors is rather common (Schäfer et al., 2016). When stressors co-occur, they can interact in complex and unexpected ways, resulting in non- additive combined effects that can be difficult to predict (Birk et al., 2020; Orr et al., 2020; Piggott et al., 2015). Indeed, the co-occurrence of two stressors can result in dominant, additive, or interactive (either synergistic or antagonistic) effects (e.g., Birk et al., 2020). The spatial arrangement of stressors along the river network can also affect their impact on running water systems. River networks form meta-ecosystems, i.e., a set of local ecosystems connected by spatial flows of energy, material, and organisms across ecosystem boundaries (Gounand et al., 2018; Loreau et al., 2003). Local stressors can alter these flows, with their effects being exported across connected habitat patches within the networks, typically with current directionality, and affect the downstream communities. A few pioneering studies have shown that local effects can propagate to downstream reaches, triggering cascading effects on communities (Chará-Serna & Richardson, 2021; Harvey et al., 2017; Negrín Dastis et al., 2024). However, the spatial component has seldom been considered in multiple stressor studies, leaving a research gap concerning the potential effects of their spatial distribution.

Flow intermittency is a growing phenomenon in running waters (Datry, Truchy, et al., 2023). The natural flow regime of many rivers and streams has been modified worldwide by the combined effects of direct anthropogenic pressures (e.g., water abstraction for agriculture or other human uses) and climate change (Larned et al., 2010), increasing the spatial and temporal extent of drying events (Sarremejane et al., 2019). It has been estimated that non-perennial rivers and streams account for more than 60% of the global river network (Messager et al., 2021) and their share is predicted to increase in the next years (Döll & Schmied, 2012). The disruption of the natural flow regime exerts a wide range of impacts on freshwater communities, like general loss of biodiversity and shifts in community composition and functionality (Burgazzi, Bolpagni, et al., 2020; Datry et al., 2014; Dudgeon et al., 2020). Flow intermittency and consequent flow reduction also lead to changes in the drift rates of macroinvertebrates, with passive drift decreasing in response to low water velocities, and active drift increasing as a stressor-avoidance mechanism (Aspin et al., 2019; Dewson et al., 2007; James et al., 2009). The spatial arrangement of intermittent and perennial sections may also affect the local effects of drying events (Datry, Boulton, et al., 2023; Leibowitz et al., 2018; Stubbington et al., 2017) through changes in drift rates resulting from alterations in connectivity or preventing recolonization processes, hampering the resistance and resilience of macroinvertebrate communities.

Light pollution is an emerging alteration of the landscape worldwide (Hölker et al., 2010; Kyba et al., 2017) particularly widespread near freshwaters (Gaston et al., 2015; Manfrin et al., 2017, 2018). Most of the in-situ studies report light pollution values in the range of 1-20 lux for freshwater systems, while natural levels reach 0.3 lux in full moon conditions (Jechow & Hölker, 2019). Moreover, recent estimates of increases in light pollution found an exponential increase, with an average growth of sky brightness of 10% per year (Falchi & Bará, 2023). Due to the perturbation of natural light cycles, light pollution can modulate both the occurrence and abundances of different taxa, for example through changes in their drift behavior (Perkin et al., 2014), reducing the drift rates of several taxa for the increased predation risk (Henn et al., 2014). For example, it may change the predatory behavior of fish on macroinvertebrates (Becker et al., 2013), potentially resulting in indirect effects downstream or on the surrounding riparian areas when the emergence and nocturnal activity of animals is changed (Thomas et al., 2016). For these reasons, light pollution can exert cascading effects on food webs, and connected ecosystems downstream or in the riparian areas (Manfrin et al., 2017).

Flow intermittency and light pollution have been seldom studied jointly (Hölker et al., 2021; Perkin et al., 2011) although they are likely to co-occur due to the growing human impacts on freshwater ecosystems. The co-occurrence of flow intermittency and light pollution could further reduce macroinvertebrate drift rates, consequently depleting downstream communities due to a lower number of exported organisms and altering sink-source dynamics in the network (Harvey et al., 2020). Conversely, the presence of unimpacted upstream habitat patches could mitigate the effect of stressors, acting as a refuge and a source of organisms and resources (Schneeweiss et al., 2022; Terui et al., 2018). Here, we present the results of a flow-through mesocosm study on the single and combined effects of flow intermittency and light pollution on macroinvertebrates (in terms of richness, abundance, and community structure) in a river network context (Figure 1). We adopted a network design aiming at evaluating the local effects of stressors and their interaction on drifting and benthic freshwater macroinvertebrates (A1), and how the effect of stressors and their spatial arrangement (single or combined) propagates along the river network, affecting downstream communities (A2). According to previous findings (e.g., Beermann et al., 2018; Datry et al., 2014; Henn et al., 2014), we expected flow intermittency to decrease taxa richness and abundance of sensitive benthic macroinvertebrates (e.g., Ephemeroptera, Plecoptera and Trichoptera) and change the drift rates depending on the dispersal ability of the organisms (HP1.1). Light pollution was expected to reduce drift rates in response to an increased predation risk. This would be mirrored by a local increase of richness and abundance in the benthic macroinvertebrate community, as light pollution represents a barrier to further dispersion (HP1.2). When the two stressors co-occur, we expected synergistic interactions with a further decrease in drift (as organism exposed to light and flow intermittency would decrease their drift disregarding the dispersal ability) coupled with decreased richness and abundance in benthos because of the effect of flow reduction on sensitive taxa (HP 1.3). Lastly, we expect that the spatial arrangement of stressors influences their effects on downstream sections. More specifically, when stressors are spatially separated in the upstream sections (i.e., they occur in different tributaries), downstream effects will be stronger, due to the lack of the buffer effect of unimpacted sections (HP2).

**Figure 1.**
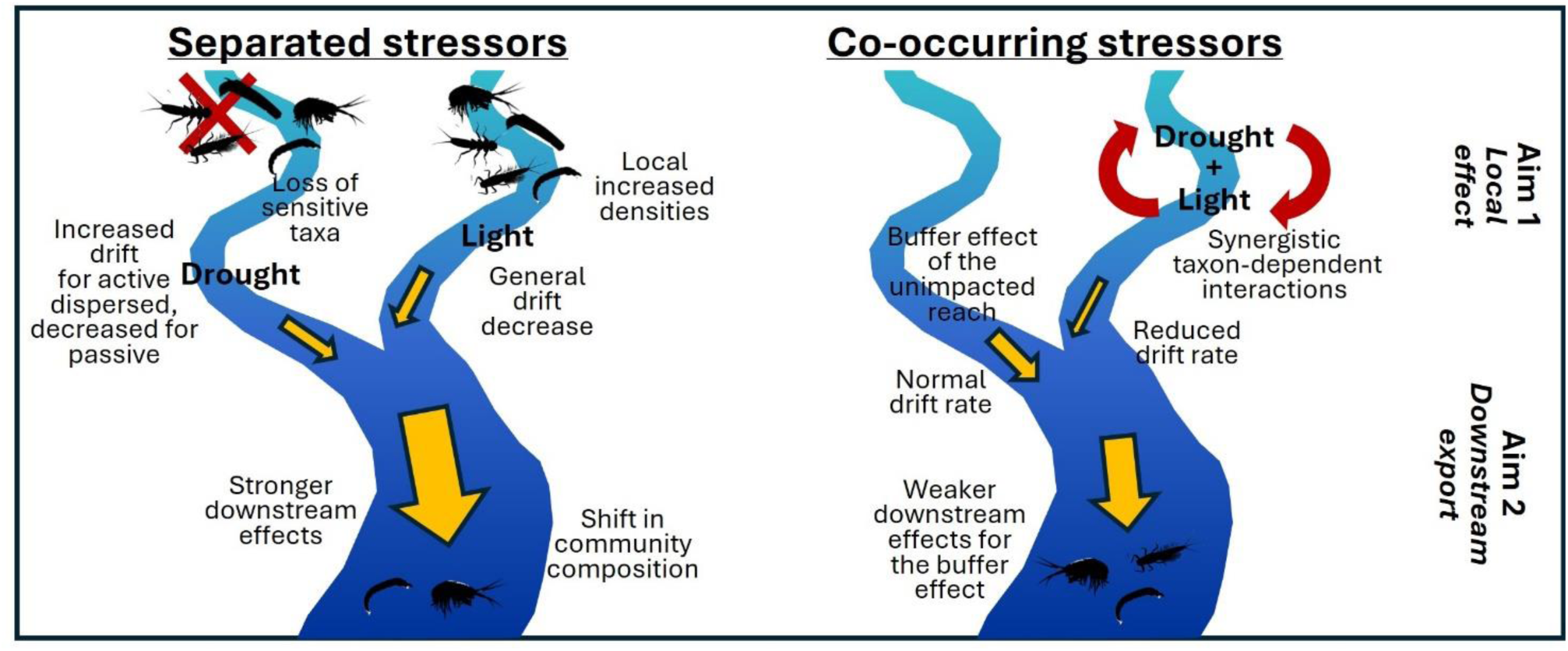
Scheme of aims and hypotheses. The first aim of this study was to assess the local effects of stressors and their interactions on macroinvertebrate communities. Under drought treatment, we hypothesized local loss of sensitive taxa and changes in export depending on the dispersal ability of organisms (HP 1.1), whereas under light treatment, we expect a general decrease in drift mirrored by increased abundances in benthos (HP 1.2). When stressors co-occur, we expect non-additive taxon-dependent effects on communities (HP 1.3). The second aim was to evaluate how the effect of stressors and their spatial arrangement (single or combined) propagates along the river network. We hypothesized stronger downstream effects, with shifts in community composition, when stressors are separated, for the lack of the buffer effect of the unimpacted reach (HP 2).

## 2. Materials and Methods

### 2.1. Experimental design and schedule

The experiment was performed in a system of 18 flow-through mesocosms (the *Disconnected* system, Figure 2a, d) at the Eußerthal Ecosystem Research Station (EERES, RPTU Kaiserslautern-Landau), in the Palatinate Forest (Rheinland-Pfalz, Germany). The EERES is located in a UNESCO biosphere reserve, densely forested and with low anthropogenic impact (Stehle et al., 2022). Each mesocosm unit (hereafter referred to as flume) was 6 m long, 14 cm wide, and 7 cm deep. The flumes were designed to simulate a simplified river network, with two separated upstream sections joining downstream. The 18 flumes were organized into six groups of three, with each upstream section independently connected to a header tank (Figure 2b). Water and drifting organisms were constantly pumped (ABK200D-2HP, Leo Group Ltd) through a 5 mm filter (placed at the water inlet) from the Sulzbach stream (a least-impacted stream running along the research station) and returned to the Sulzbach downstream of the system. Discharge was kept constant (with daily calibration) at 36 L/min per flume, resulting in flow velocity and water depth ranging from 22 cm/s and 4 cm to 17 cm/s and 7 cm (in upstream and downstream sections respectively, Figure S1). Each flume was supplied with a mix of sediment: 9.1 L of gravel (Ø 0.2-2 cm; 70% of substrate), 1.3 L of sand (Ø < 0.2 cm; 10% of substrate), 1.3 L of cobbles (Ø 2-6 cm; 10% of substrate) and 1.3 L of stones (Ø > 6 cm; 10% of substrate). This substrate composition reflected the composition in the Sulzbach stream. Throughout the experiment, we monitored physical (conductivity, dissolved oxygen, pH, and water temperature, with multiparametric probe WTW Multi 3630 IDS Set G) and hydraulic (flow velocity and water depth, with an OTT MF pro electromagnetic current meter) parameters weekly (Figure S1).

**Figure 2.**
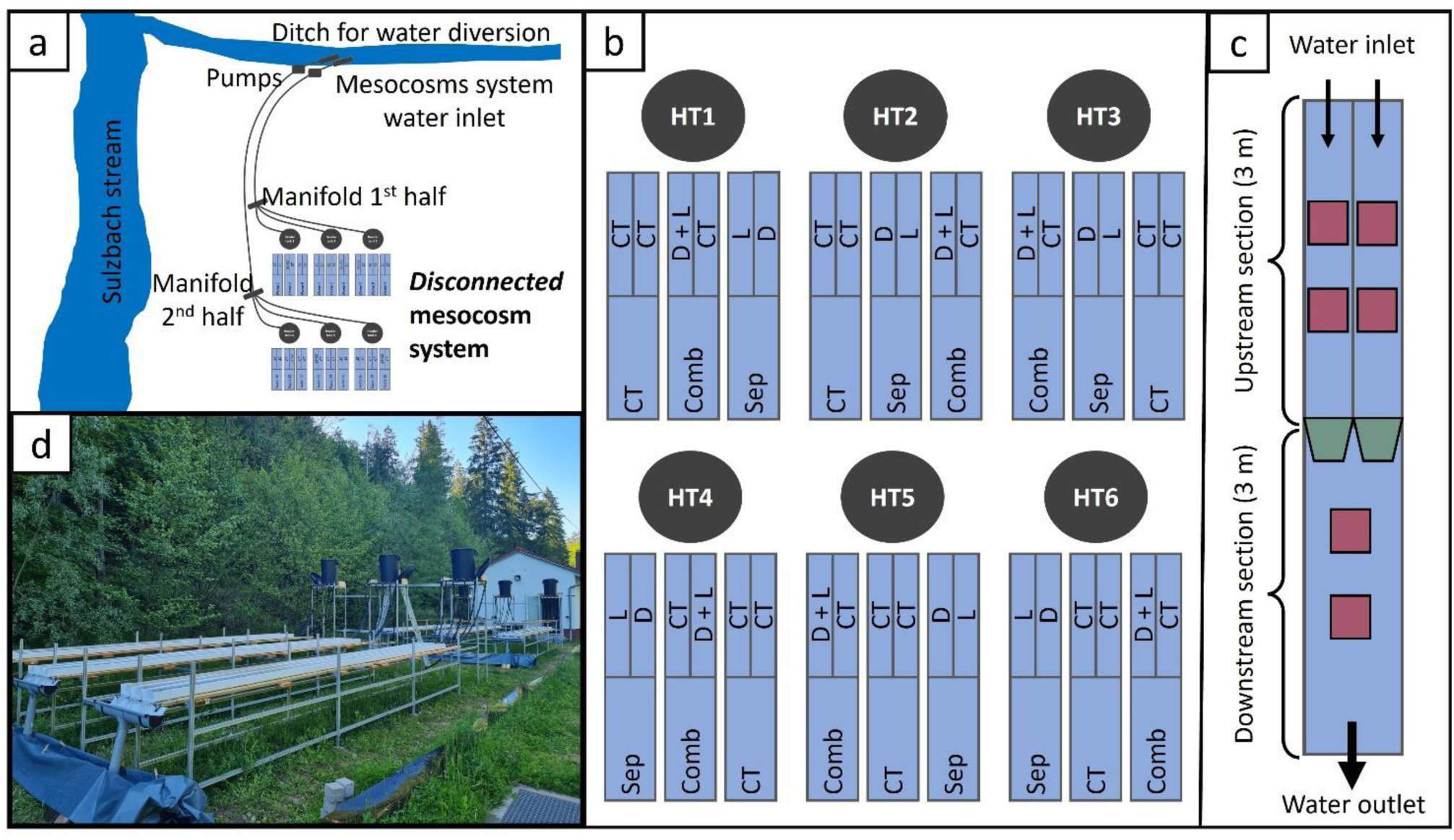
Scheme of the *Disconnected* mesocosm system (a). Experimental design (HT = header tank) and randomized block allocation of the stressors (D = drought treatment, L= light treatment, CT = control, Sep = separated upstream stressors, Comb = combined upstream stressors) in the upstream sections (b). Scheme of the sampling spots in each flume (c), with red areas for benthos T1, and green areas for the position of the drift nets. Picture of the mesocosm system (d).

The flumes were initially passively colonized for approx. three weeks (Table S1), starting from 06^th^ July 2021 (day -26). Before applying the treatments (day -9), we supplied the flumes with an active seeding of macroinvertebrates. Organisms were collected in the Sulzbach stream, with 3-mins kick-sampling in a total area of 18 m^2^ (sampling approx. 1 m^2^ per flume). After sampling, organisms were homogeneously distributed in the flumes (both in upstream and downstream sections), briefly stopping the water flow (for 3 minutes) to allow macroinvertebrates to settle. On 01^st^ August 2021 (day 0), we started the treatment phase. Stressors (flow intermittency and light pollution) were applied with a randomized block design in the upstream sections, resulting in one control flume (i.e., with both upstream sections being unimpacted), one flume with separate stressors (i.e., with flow intermittency applied in one upstream section and light pollution in the other one), and one flume with co-occurring stressors (i.e., with flow intermittency and light pollution being applied in the same upstream section, leaving the second one unimpacted) for each header tank (Figure 2b). For the flow intermittency treatments (hereafter referred to as *drought* treatment), taps feeding the upstream sections were closed, simulating the ponded phase of the drying process (Hill & Milner, 2018). This translated into flow velocity reduced to zero in drought treatment sections, and discharge halved in all downstream sections of treatment flumes. For light pollution treatments (hereafter referred to as *light* treatment) LED strips (IP65300 SMD 2835,12 V, 4000K, Yphix) were placed 40 cm above the water surface (running all along the upstream sections) and calibrated weekly to 10 lux at the benthos level (MW 700 Portable LuxMeter, measured keeping the sensor submerged at the bottom of the water column). The LED strips were connected to timers and automated to turn on at 9 pm and off at 6 am. To prevent cross-contamination in sections without light treatment, we installed black fabric panels around the light treatment sections and measured light intensity on the other side of the black fabric to check their efficiency (resulting in mean and standard deviation 0 ± 0 Lux for all measurements in light-free sections).

### 2.2. Sampling and lab activities

We sampled drifting macroinvertebrates weekly during the treatment phase (T1-T4, Table S1). Drift nets (mesh size 0.5 mm) were placed at the outlet of each upstream section for 24 hours, to collect integrated day and night drift samples. Benthic macroinvertebrates were collected at the end of the treatment phase (day 22, Table S1), sampling areas of 150 cm^2^ with two replicates in each section (Figure 2c). Macroinvertebrate samples (both for drift and benthos) were preserved in 96% ethanol. Organisms were counted and identified to a mixed taxonomic level (family or genus) following Tachet et al. (2010). After counting and identifying, we obtained the multivariate community matrices and computed the metrics taxa richness (i.e., sum of all taxa) and abundance (i.e., sum of all organisms) of each sample, both for benthos and drift. All taxa were categorized as active or passive dispersers by combining different drift propensities (Sarremejane et al., 2020), dispersal strategies, and locomotion traits (Tachet et al., 2010). See Table S2 in the supplementary for more details.

### 2.3. Data analysis

We assessed the local effect of single and combined stressors on community metrics (richness and abundance) with mixed modeling, both for drifting and benthic macroinvertebrates. We coded drought and light treatments as dummy variables and used them and their interaction as fixed factors. In the random part of the models, we included the nested structure header tank/flume for benthos and drift. For the drift dataset, we also considered time (categorical; T1, T2, T3, and T4) as a fixed factor. To assess the downstream propagation of the effects of the stressors on community metrics, we run mixed models using the upstream arrangement of the stressors (categorical; control, separated, and combined) as the fixed factor (plus time for the drift dataset), and flume nested within header tank as the random factor. For the drift dataset, we summed up the data from the two sides within each flume, to obtain the total downstream export. We summarized the results of the models with Type II ANOVA (to account for the unbalanced design of the control treatment in the tributaries) and used the marginal and conditional R^2^ to assess their goodness of fit. For downstream models when treatment was significant, we conducted pairwise Least Squares-means post-hoc tests with Bonferroni adjustment to detect the differences between combined and separated.

To assess the local and downstream effects of single and combined stressors on community composition, we applied multivariate generalized linear latent variable models (GLLVMs, Niku et al., 2019) to abundance data. This approach represents a multivariate extension of generalized linear models, which allows the assessment of the joint response of all taxa to environmental gradients (Niku et al., 2019; Pollock et al., 2014). GLLVMs follow the basic form:

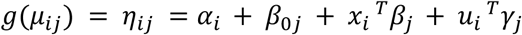

Where *g*() is a link function, *η*_*i,j*_ a linear predictor for the site *i* and taxon *j*, *a*_*i*_ a site-specific intercept, *β*_0*j*_ a species-specific intercept, *x*_*i*_ are environmental covariates, T marks the transpose, and *β*_*j*_ the corresponding regression coefficients. *u*_*i*_ are latent variables at site *i* and *γ*_*j*_ the factor loadings, which indicate how species *j* responds to latent variables.

Separate GLLVMs were run for drift upstream, benthos upstream, drift downstream, and benthos downstream. We only retained taxa with abundance > 0.5% of total abundance for each dataset (Table S2). For the upstream drift and benthos data, we tested the effect of drought and light (and their interaction) coded as dummy variables and included the same nested structure (header tank/flume) as for the univariate metrics models. These two upstream models were coded with five latent variables. For the downstream data, we tested the effect of treatment (being a factor with three levels: control, separated, and combined), also including the flume nested within the header tank as random effect. In this case, we fit the two downstream models with three latent variables. Both for upstream and downstream models the number of latent variables was optimized based on the Akaike information criterion (AIC). In drift models, both upstream and downstream, we also included time as a fixed factor. For all GLLVMs, we used Gaussian variational approximation method (Hui et al., 2017), and assumed a Poisson distribution of the response variables. All models were checked for overdispersion with diagnostic plots. We assessed the amount of variation in the data explained by the stressors for each model using the trace of the residual covariance matrix as a measure of unexplained variation. In GLLVMs, the residual covariance matrix *Σ* can be obtained as *Σ* = *ΓΓ*^*T*^, where Γ is the matrix of factor loadings. We then compared this quantity before and after stressors were included in the model (i.e., comparing the unconstrained and constrained models, Niku et al., 2019). This was done according to the following equation:

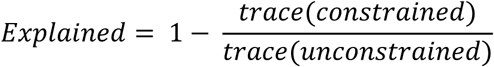

All analyses (significant threshold: p = 0.05) were performed with the packages *dplyr* (version 1.1.0; Wickham et al., 2023), *tibble* (version 3.1.8; Müller et al., 2023), *tidyr* (version 1.3.0; Wickham et al., 2024), *lme4* (version 1.1-31; Bates et al., 2019), *ggplot2* (version 2_3.5.0; Wickham et al., 2019), *ggtext* (version 0.1.2; Wilke & Wiernik, 2022)*, vegan* (Oksanen et al., 2019), *gllvm* (Niku et al., 2019), *car* (version 3.1-1; Fox & Weisberg, 2018), *MuMIn* (version 1.46.0; Bartoń, 2023), *lsmeans* (version 2.30-0; Lenth, 2016), *forcats* (version 1.0.0; Wickham, 2023), and *patchwork* (version 1.2.0; Pedersen, 2024) of the R statistical software (R version 4.1.2, R Core Team, 2021). The metrics values have been log-transformed when needed for mixed models. All data from this study are openly available on Dryad and the R code can be found at https://github.com/GSBurg/Disconnected. A roadmap of the data analysis is reported in Supplementary materials (Table S4).

## 3. Results

During the colonization phase, all environmental parameters (Figure S1) were comparable among flumes, with a difference between upstream and downstream sections only for flow velocity, which decreased downstream by 20%. During the treatment phase, flow velocity was null in drought treatment sections, with associated increases in dissolved oxygen, water temperature, and pH and stronger variability in conductivity. Flow velocity also decreased downstream in all treatment flumes, with a consequent decrease in water depth due to sediment deposition. Light treatment had no clear effect on environmental parameters, besides a slight increase of the dissolved oxygen during the daytime (on average, from 10.22 mg/L in control sections to 10.30 mg/L in the light treatment section).

In total, we collected 5564 drifting and 11326 benthic macroinvertebrates, belonging to 39 and 40 taxa respectively. Among these, 30 taxa were found in both drift and benthos samples, whereas 19 taxa were either exclusive of drift or benthos (Table S3). Both drift and benthos were dominated by few, highly abundant taxa (Table S3) namely the Diptera Orthocladiinae, Chironomini, and Tanypodinae, and the two genera *Baetis* sp. (Order Ephemeroptera) and *Gammarus* sp. (order Amphipoda). The rest of the taxa composing the community of drift and benthos had a relative abundance lower than 5%.

### 3.1. Upstream local effect of single and combined stressors on drifting and benthic macroinvertebrates

Regarding the local effects of single and combined stressors, we expected drought treatment to have a negative effect on both benthos and drift (but with differences depending on dispersal abilities), light pollution to reduce macroinvertebrate drift rates and the combination of the two stressors to further reduce drift and benthos due to synergistic interactions.

For the drift, we found a significant effect of drought and light treatments, as well as their interaction for taxa richness, while for abundance we found a significant effect of drought treatment and the interaction of drought and light, but no effect of the light treatment alone (Table 1, Figure S2). On average, drought treatment reduced taxa richness and abundance by 50 and 70% respectively, whereas richness increased by 10% under light treatment alone. The co-occurrence of stressors led to a weak synergistic interaction, with a lower richness and abundance than the additive effects of single stressors (Figure S2). The richness and abundance of drifting macroinvertebrates increased significantly over time.

**Table 1.** Mixed models result for upstream sections of drift and benthos data, summarized with type 2 ANOVA. Significant results are marked in bold. R^2^m = marginal R^2^ (i.e. variance explained by the fixed effects); R^2^c = conditional R^2^ (i.e. variance explained by the entire model); Chisq = chi-square values; Df = degrees of freedom; HT = header tank.

| Response variables | R<sup>2</sup>m | R<sup>2</sup>c | Explanatory variables | Chisq | Df | p-value |  |
| --- | --- | --- | --- | --- | --- | --- | --- |
| DRIFT - Model structure: drought* light + time + (1\|HT/flume) |  |  |  |  |  |  |  |
| Taxa richness | 0.638 | 0.648 | Drought | 229.74 | 1 | <0.001 | *** |
|  |  |  | Light | 6.59 | 1 | 0.010 | * |
|  |  |  | Time | 15.48 | 3 | 0.001 | ** |
|  |  |  | Drought*Light | 6.42 | 1 | 0.011 | * |
| Abundance | 0.692 | 0.707 | Drought | 299.41 | 1 | <0.001 | *** |
|  |  |  | Light | 0.93 | 1 | 0.336 |  |
|  |  |  | Time | 9.50 | 3 | 0.023 | * |
|  |  |  | Drought*Light | 4.01 | 1 | 0.045 | * |
| BENTHOS - Model structure: drought* light + (1\|HT/flume) |  |  |  |  |  |  |  |
| Taxa richness | 0.333 | 0.426 | Drought | 24.36 | 1 | <0.001 | *** |
|  |  |  | Light | 5.09 | 1 | 0.024 | * |
|  |  |  | Drought*Light | 0.29 | 1 | 0.593 |  |
| Abundance | 0.420 | 0.549 | Drought | 45.32 | 1 | <0.001 | *** |
|  |  |  | Light | 2.00 | 1 | 0.157 |  |
|  |  |  | Drought*Light | 1.01 | 1 | 0.315 |  |

For the benthos, light alone significantly affected taxa richness only, with an average reduction of 17%, whereas in drought treatments, we observed a reduction of both richness and abundance by 33 and 64% respectively. We detected a dominance of drought treatment over light, without significant interactions among stressors (Table 1, Figure S2).

Stressors explained 70% and 41% of the variation in drift and benthos communities, respectively. The local effect of the stressors differed depending on their spatial arrangement, with taxon-specific changes (Figure 3). For most taxa, drift decreased under drought treatment and increased under light treatment, with differences depending on the dispersal ability of the organisms only in light treatments, where we observed increased drift mainly (but not only) for active dispersers. In benthic communities, drought treatment reduced the abundance of most taxa, whereas light treatment only affected the dipterans Tanytarsini. We found significant antagonistic or synergistic interactions for three drifting (*Polycelis* sp., *Baetis* sp., and *Ephemerella* sp.) and one benthic taxon (Simuliidae). The abundance of drifting organisms increased significantly over time.

**Figure 3.**
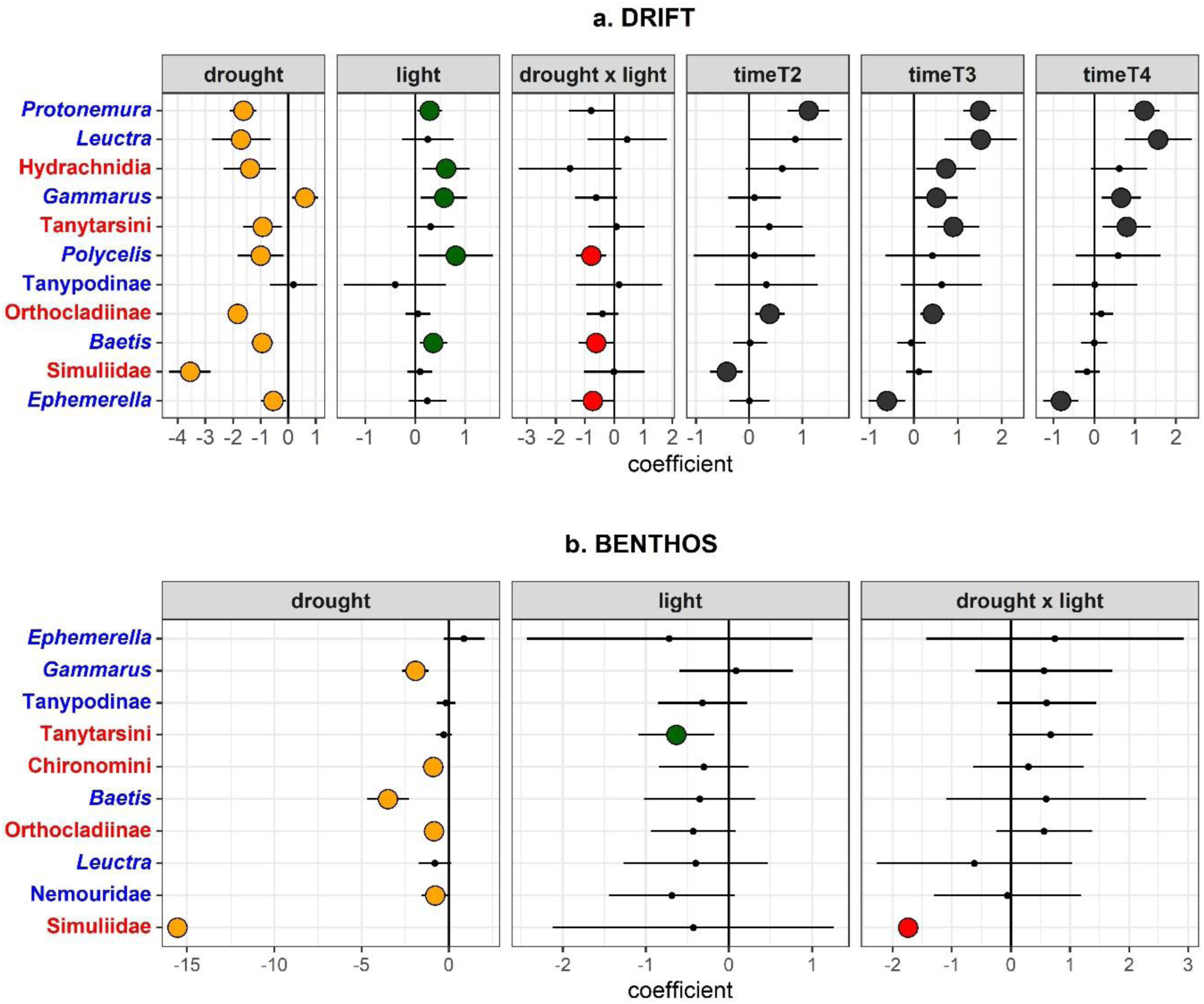
Coefficient plots from generalized linear latent variable models (GLLVMs) for drifting (a) and benthic (b) macroinvertebrates from upstream sections, showing the effect of stressors and their interaction (and time for drift data) on the most common taxa (being taxa with abundance > 0.5%). *Elmis* sp. and Naididae were removed from drift upstream plots because of the excessively large error bars. Labels of taxa names are written in blue for active dispersers and in red for passive dispersers.

### 3.2. Downstream export of the effects of the single and combined stressors on drifting and benthic macroinvertebrates

We expected a downstream spatial cascading effect on macroinvertebrate communities, depending on the upstream spatial arrangement of stressors (single or combined), with stronger downstream effects for drought and light treatments applied separately in upstream sections, due to the lack of the buffer effect of the unimpacted section.

According to our results, upstream stressors and their spatial arrangement significantly affected downstream communities. Upstream treatments significantly affected the abundance of drifting macroinvertebrates, with the export decreasing over time from control over separated to the combined treatments (Table 2, Figure S3). This reduced export, however, was not mirrored by changes in richness or abundance in downstream benthos (Table 2), for which we only found a non-significant trend (Figure S3).

**Table 2.** Mixed models result for downstream sections of drift and benthos data, summarized with type 2 ANOVA. Significant results are marked in bold. R^2^m = marginal R^2^ (i.e. variance explained by the fixed effects); R^2^c = conditional R^2^ (i.e. variance explained by the entire model); Chisq = chi-square values; Df = degrees of freedom; HT = header tank.

| Response variables | R<sup>2</sup>m | R<sup>2</sup>c | Explanatory variables | Chisq | Df | p-value |  | Treatment differences |
| --- | --- | --- | --- | --- | --- | --- | --- | --- |
| DRIFT - Model structure: treatment + time + (1\|HT/flume) |  |  |  |  |  |  |  |  |
| Taxa richness | 0.212 | 0.254 | treatment | 3.14 | 2 | 0.208 |  | none |
|  |  |  | time | 16.28 | 3 | 0.001 | *** |  |
| Abundance | 0.361 | 0.460 | treatment | 34.09 | 2 | <0.001 | *** | all |
|  |  |  | time | 13.45 | 3 | 0.004 | ** |  |
| BENTHOS - Model structure: treatment + (1\|HT/flume) |  |  |  |  |  |  |  |  |
| Taxa richness | 0.070 | 0.258 | treatment | 2.20 | 2 | 0.333 |  | none |
| Abundance | 0.152 | 0.413 | treatment | 4.78 | 2 | 0.092 |  | none |

Stressors upstream arrangement explained 27% and 3% of the variation in downstream drift and benthos community composition, respectively (Figure 4). Most of the taxa showed a stronger decrease in abundance in combined treatments (e.g. Orthocladiinae, Simuliidae, and *Ephemerella* sp. in drift, Simuliidae, *Baetis* sp., and *Gammarus* sp. in benthos). Only in two cases, we detected downstream increases: *Gammarus* sp. in drift communities and Orthocladiinae in benthos communities. For drifting macroinvertebrates, few taxa (*Ephemerella* sp., *Protonemura* sp., and *Elmis* sp.) responded only to the combined treatments.

**Figure 4.**
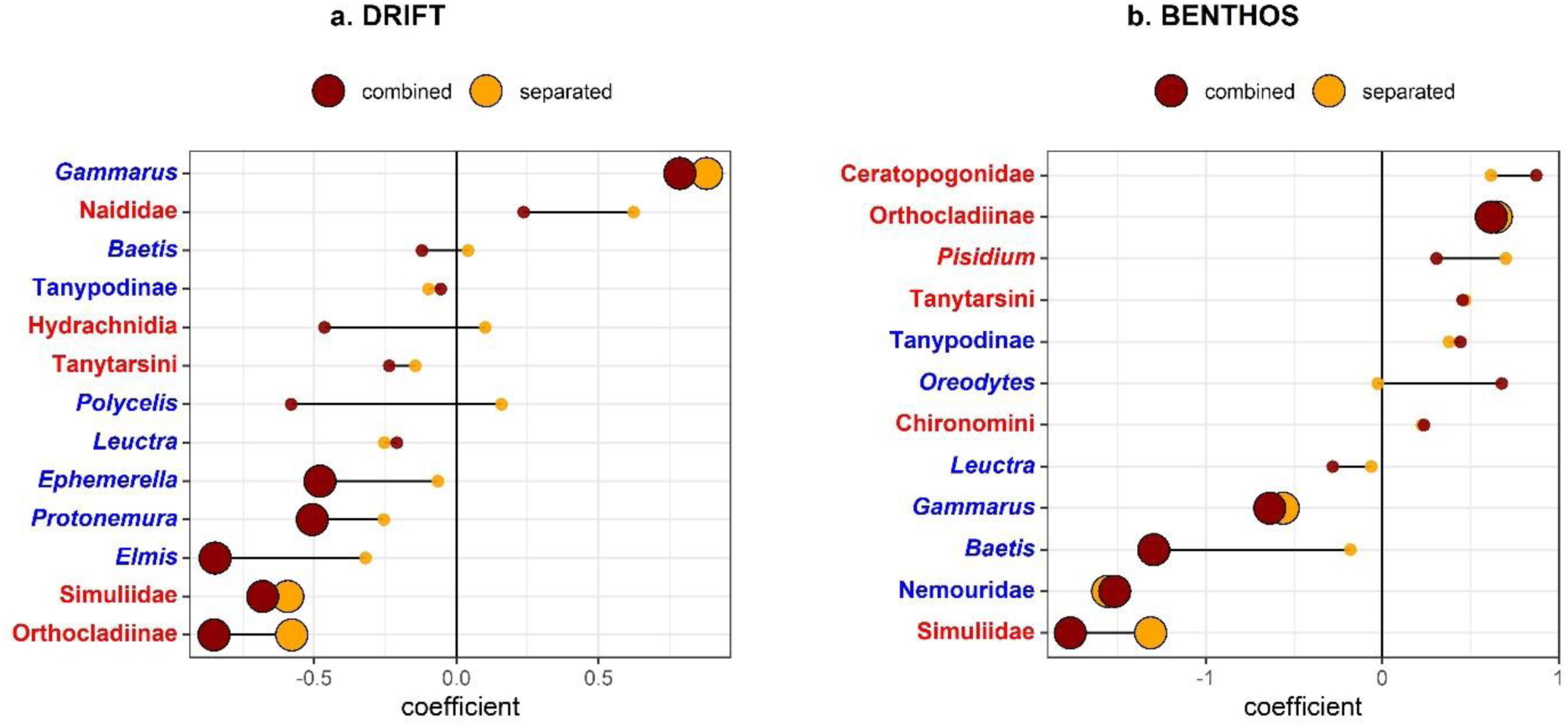
Coefficient plots from generalized linear latent variable models (GLLVMs) showing the effect of treatment arrangement (combined or separated) on downstream drifting (a) and benthic (b) communities (considering only common taxa, having abundance > 0.5%). Red dots represent combined treatments, and yellow dots separated treatments. Big dots represent a significant effect of treatment arrangement on downstream communities, whereas small dots are for non-significant effects. Labels of taxa names are written in blue for active dispersers and in red for passive dispersers.

## 4. Discussion

With the growing human presence, multiple stressors are spreading in river networks, representing a challenge for biodiversity conservation, as their potential interactions are often difficult to forecast and can propagate downstream. Here, we assessed the single and combined effects of flow intermittency and light pollution on macroinvertebrate communities, both locally, and in downstream unimpacted reaches. Results suggest that stressors and their interactions affect macroinvertebrates locally, with either dominance of drought treatment or interactive effects between drought and light. Moreover, the effects propagate along the river network, affecting downstream communities with more severe impacts when stressors co-occur upstream.

### 4.1. Upstream local effect of the single and combined stressors on drifting and benthic macroinvertebrates

In our study, we found different effects of the stressors on the drifting and benthic macroinvertebrates, often with metric- and taxa-specific responses. Contrasting treatment effects were found in the drift, where richness and abundance decreased in drought treatments and increased in light treatments, while their co-occurrence led to local interactions. For benthic macroinvertebrates, both stressors decreased the values of richness and abundance, but with stronger and mainly dominant effects of drought treatment.

Drought treatment had in general a negative effect, both on drift and benthos, partially supporting our hypothesis (HP 1.1). Previous findings from studies about drift rates are contrasting, reporting both increases (González et al., 2018; James et al., 2009) and decreases (Calapez et al., 2017) of the drift rates under flow intermittency. This difference can be explained by differences in drought intensity or duration, or by a different ratio of active and passive dispersers in the communities (Dewson et al., 2007), which depends on the local species pool in different biogeographic areas. However, in our study, the effects of flow reduction stressor did not differ between active and passive dispersers. For example, when examining taxon-specific changes in drift under drought treatment (Figure 3a), we found a decreased drift for five active dispersers (*Protonemura* sp., *Leuctra* sp., *Polycelis* sp., *Baetis* sp., and *Ephemerella* sp.) and four passive dispersers (Hydrachnidia, Tanytarsini, Orthocladiinae, and Simuliidae). The observed decreased drift was associated with reduced values of richness and abundance in benthos, with values halved compared to the controls. These results highlight how flow intermittency and associated flow reduction can strongly affect macroinvertebrate communities in temperate areas even in the early stages, like the ponded phase of the drying process (Chadd et al., 2017). The transition from normal flow to ponded phase implies changes in several parameters, like flow velocity, oxygen availability, and water temperature. This happened also in our study system, with all the background environmental parameters (flow velocity, water depth, dissolved oxygen, temperature, conductivity, and pH) altered in the tributaries where we applied the drought treatment (Figure S1). Organisms with narrow niches cannot tolerate these changes and they usually either escape from the stressor (via drift, emergence or using in-stream refuges like the hyporheic zone) or die (de la Fuente et al., 2018). In our experiment, the reduction in benthos abundance was not mirrored by an increased drift under drought treatments, suggesting that the organisms either did not actuate the avoiding strategy and died in the system or accelerated their emergence (Hunn et al., 2024). While emergence rate was not measured in this study, the lack of adaptation of the local species pool (macroinvertebrates in the system were from a pristine perennial stream), the way we applied the stressor (usually, flow intermittency is a ramp disturbance, gradually increasing, while in our experiment it was suddenly applied) or the lack of connection with the hyporheic zone (due to how we build the flumes) could explain this finding (Bruno et al., 2020; Datry, Truchy, et al., 2023; Lake, 2000).

Contrary to our expectations (HP 1.2), light pollution increased the taxon-specific drift rates of macroinvertebrates and overall taxa richness in drift. Most of the previous studies report decreases in drift rates in light pollution conditions (e.g., Henn et al., 2014; Perkin et al., 2014), which is usually attributed to predation avoidance behavior for the increased predation risk. The lack of large predators like fish and other vertebrates in our system may explain this discrepancy. While drift-feeding fishes visually search for drifting prey (Neuswanger et al., 2014), macroinvertebrate predators detect their prey by both visual and mechanical cues (Peckarsky, 1982). In the latter case, increased nocturnal light would only have a minor effect on predators behavior and predation risk. Consequently, macroinvertebrates would react to light treatments with avoidance mechanisms, increasing their drift rate. The increased drift rates in light treatments found in our experiment could also be an indirect consequence of light pollution. Previous studies reported a decrease in biomass and compositional shifts in periphyton exposed to LED light treatment (Grubisic et al., 2017; Grubisic, Singer, et al., 2018; Grubisic, van Grunsven, et al., 2018). Reduced biomass at the base of the food web would induce bottom-up effects on grazing macroinvertebrates, depleting their food resources and driving them to drift in search of food. In a companion study (Juvigny-Khenafou et al., 2024), light treatment increased the abundance of macroinvertebrates in leaf bags, but the leaf decomposition rate remained similar. This suggests that leaf litter consumption did not change and that litter was primarily used by some benthic macroinvertebrates as an in-stream refuge (Burgazzi, Laini, et al., 2020), thus the increased drift rate we found for several taxa might be induced by research for food. As we did not find major taxon-specific responses of benthic common taxa under light treatments, but taxa richness decreased in benthos and increased in drift, it is possible that also rare taxa significantly contributed to increasing the overall drift richness. The only taxon affected by light treatment in benthos was Tanytarsini (family Chironomidae), which significantly decreased. While dispersal ability or feeding habits alone do not explain this finding, their lower benthic abundance could be a consequence of their smaller size (Schmidt-Kloiber & Hering, 2015), which together with limited mobility makes them easy prey for other macroinvertebrates.

According to our results, drought and light treatments significantly interacted, leading to metric and taxon- specific effects, especially for drifting macroinvertebrates. Contrary to our expectation (HP 1.3), the observed significant interactions were both antagonistic and synergistic (e.g., taxon-specific effects in drifting and benthic communities), with resultant effects either in the middle or stronger than single ones. For benthic macroinvertebrates, however, we mainly found a dominance of drought treatment over light treatments, most likely due to the several changes in environmental variables consequent to the induced ponded phase. To our knowledge, our experiment is the first study testing the combined effect of these two stressors. More targeted research would be needed to clarify the mechanisms underlying these interactions. However, from what we observed in our system it seems that in most cases macroinvertebrates show opposite reactions to flow reduction and light pollution, with stronger effects of flow reduction, which are in some cases attenuated or nullified by the presence of light via additive or antagonistic interactions.

### 4.2. Downstream export of the effects of the single and combined stressors on drifting and benthic macroinvertebrates

Multiple stressors are seldom studied in a spatial context, with their potential cascading effects in downstream unimpacted reaches still quite unknown. A recent study tested whether sediment and nutrient enrichment in tributaries affect downstream ecosystems (Chará-Serna & Richardson, 2021). They found lower macroinvertebrate richness and density values when stressors were combined in upstream tributaries, highlighting that the effects of stressors and their interaction are exported along the river network. Similarly, our results suggest a downstream propagation of flow reduction and light pollution interaction, where local stressor arrangement influenced their effects at the river network level.

For drifting macroinvertebrates, we found differences among treatments, with total drift abundance decreasing from control to separated stressors and being lower for combined stressors (Table 2, Figure S3). We also found taxon-specific changes in community composition, with drift rates of several common taxa being lower for combined treatments (Figure 4). This suggests that stressors interact locally, and their interaction triggers cascading effects downstream via changes in the export of some organisms. The observed downstream effects were stronger when stressors co-occurred in upstream tributaries, with no buffer effect of the unimpacted tributary (Schäfer et al., 2017), contradicting our initial hypothesis of stronger downstream effects for separated upstream stressors (HP 2). Moreover, this greater decreased drift in combined treatment does not seem to be related to dispersal ability, as both active and passive dispersers are equally affected. Such changes may have severe consequences at the river network level, through changes in meta-communities and meta-ecosystem dynamics (Harvey et al., 2020; Naman et al., 2016). In fact, a reduced drift from upstream patches can lead to food scarcity for fish communities or to alterations in primary producers consumption and organic matter degradation, with potential cascading effects on the food web (Wipfli & Baxter, 2010). For example, in our study, we found reduced downstream export in flumes with combined stressors for the taxa *Elmis* sp., *Ephemerella* sp., and Orthocladiinae. These taxa are scrapers, i.e., they feed on the biofilm growing on available submerged substrates in running waters, grazing autochthonous primary producers. Without the top-down control of scrapers, primary producers in downstream sections may rapidly proliferate up to nuisance levels (Sturt et al., 2011), with further negative impacts on ecosystems (Burgazzi, Bolpagni, et al., 2020). Our study showed that the downstream consumption of allochthonous organic matter could be affected as well, for example, due to the decreased export of the shredder taxa like *Protonemura* sp.. Feeding on allochthonous organic matter, shredders both support the detritus chain in river networks and reduce the clogging of riverbeds with decomposing organic matter (Suter II & Cormier, 2015). This highlights that multiple stressors arrangement and their downstream propagation not only affect taxonomic composition of macroinvertebrate communities but can also lead to functional alteration, compromising an ecosystem equilibrium.

The difference in downstream export was not mirrored by significant changes in the overall benthic community downstream. Taxa-specific effects were found for combined and separated treatments, but the share of explained variance by the downstream benthos model was however low. This suggested that organisms drifted out of the system, without settling in downstream benthic habitats. Drought treatment, which was the dominant stressor in several cases, also induced changes in environmental parameters downstream, halving the flow velocity and increasing fine sediment deposition with consequent depth reduction and substrate clogging (Bendaoud et al., 2021; Mathers et al., 2022). All these factors may contribute to making downstream habitats less suitable for macroinvertebrates in our system. It is also possible that the duration of the experiment was insufficient to have detectable effects on benthic macroinvertebrates. In fact, while the drifting response can be fast (Calapez et al., 2017), shifts in benthic community composition might require a longer time, as the way stressors affect target communities depends on the nature of the stressors themselves (acting as a pulse, press or ramp; Lake, 2000), and responses are often characterized by threshold disturbance levels (Chen et al., 2023).

### 4.3. Limitations of the study and future perspectives

Mesocosms are a valuable alternative to field studies, as they allow to manipulate stressors, replicate, and reduce the noise due to other confounding factors affecting natural systems while maintaining nearly realistic environmental conditions (Schneeweiss et al., 2023). The *Disconnected* mesocosm system is one of the first ones designed to represent a simplified river network. However, to increase replication and contain the costs, we had to reduce the size of the single flumes. This might have biased our results, as on a short distance, organisms usually considered passive dispersers may be able to move actively. Also, the lack of vertical connectivity and a proper hyporheic zone might affect the response of some animals to stressors. For example, instead of drifting out of the system some of them could hide in the sediment, using the hyporheic zone as an in-stream refuge (Stubbington, 2012). The size of the flumes in the *Disconnected* system also precluded the presence of big predators such as fish, which could modify the effects of light pollution. However, the species pool in the flumes was representative of the one in the Sulzbach stream. The network design of the *Disconnected* system represents the starting point for future research, tackling the effect of different stressors, including their temporal variability or being replicated in different biogeographic regions.

## 5. Conclusions

Multiple stressor studies are quite common, but only a few of them are framed within a spatial context. Our study is one of the first contributing to a spatial perspective in multiple stressors research in freshwater ecosystems, assessing both local and downstream single and combined effects of stressors in a simplified meta- ecosystem approach. Also, despite the growing research investigating flow intermittency and light pollution, their combined effect had never been studied before. According to our result, both flow reduction and light pollution affect drifting and benthic macroinvertebrates, with their combined effects being either dominant or interactive. Moreover, we showed that the effects of these stressors and their interactions are exported along the river network, with the magnitude of these effects depending on the spatial arrangement of the stressors themselves. This claims for a change of paradigm in multiple stressor studies, not only focusing on local effects and interaction but also exploring the implications at the meta-ecosystem level. Considering the growing number of human-induced stressors and the pressing need to develop efficient conservation strategies for freshwater ecosystems, our findings represent a valuable step forward, informing our understanding of river ecosystems, not as single, disconnected, units but as complex, connected networks.

## Acknowledgments

This work is part of the DFG (Deutsche Forschungsgemeinschaft) funded Disconnected project (451144087). Alessandro Manfrin was supported by the DFG – 326210499/GRK2360. Special thanks to Tobias Graf and the EERES staff, Dr. Alexander Feckler, Dr. Lorenzo Rovelli, Helena Bayat, Stephan Kunz, Maxime Bonin, and Nikita Steiner for the support during the construction of the *Disconnected* system and the sampling campaigns.

## Author contributions

NJK, JP, EH, AT, FL, and RBS conceived the idea. GB, NJK, and RBS planned the experiment. GB and NJK built the mesocosm system and performed the experiment. GB processed the samples. GB, VCS, JJ, and AM analyzed the data. GB led the writing of the manuscript with input from all authors. All authors have approved the submitted version.

## Data availability statement

The data that support the findings of this study are openly available on Dryad. R code used for this study can be found at https://github.com/GSBurg/Disconnected.

