## Supplementary materials for "Multiple stressors in river networks: local and downstream effects on freshwater macroinvertebrates"

### List of Contents:

**Table S1**. Detailed schedule of the experiment conducted in 2021, including the activities of both colonization and treatment phases.

**Table S2**. Common taxa (abundance > 0.5%) used in generalized linear latent variable models (GLLVMs).

**Table S3**. Full taxa list with abundances for drifting and benthic macroinvertebrates.

**Table S4.** Roadmap of the data analysis, including their matching aim and the target variable/community.

**Figure S1.** Boxplots summarizing the background data (flow meter, depth, and multiparametric probe) for each treatment level during both the colonization and treatment phases.

**Figure S2**. Variability of taxa richness and abundance metrics for drifting and benthic macroinvertebrates in upstream sections.

**Figure S3.** Variability of taxa richness and abundance metrics for drifting and benthic macroinvertebrates in downstream sections.

**Table S1**. Detailed schedule of the experiment conducted in 2021, including a description of the activities of both colonization and treatment phases.

| | Date | Day | Activity | Description |
|---|---|---|---|---|
| COLONIZATION PHASE | 06 Jul | -26 | Beginning of the colonization phase | Water started to circulate in the system. Discharge set at 36 L/min per flume and kept constant with daily calibration. |
| | 08 Jul | -24 | Physical parameters | Conductivity, dissolved oxygen, pH, and temperature were measured in both upstream and downstream sections of each flume (background data, Figure S1). |
| | 09 Jul | -23 | Hydraulic parameters | Flow velocity and water depth were measured in both upstream and downstream sections of each flume (background data, Figure S1). |
| | 16 Jul | -16 | Physical parameters | Conductivity, dissolved oxygen, pH, and temperature were measured in both upstream and downstream sections of each flume (background data, Figure S1). |
| | 17 Jul | -15 | Hydraulic parameters | Flow velocity and water depth were measured in both upstream and downstream sections of each flume (background data, Figure S1). |
| | 22 Jul | -10 | Physical parameters | Conductivity, dissolved oxygen, pH, and temperature were measured in both upstream and downstream sections of each flume (background data, Figure S1). |
| | 23 Jul | -9 | Active seeding of the flumes | 3-mins kick-sampling in the Sulzbach stream. Macroinvertebrates homogenized and spread in each flume. See the methods section for more details. |
| | 24 Jul | -8 | Hydraulic parameters | Flow velocity and water depth were measured in both upstream and downstream sections of each flume (background data, Figure S1). |
| TREATMENT PHASE | 01 Aug | 0 | Beginning of the treatment phase + drift nets placed | Taps closed in the sections with flow intermittency treatment (drought and drought + light) for simulating the ponded phase. LED strips were placed over the sections with ALAN treatment (light and drought + light). Black fabric barriers were installed to prevent cross-contamination. Drift nets were placed at the outlet of each upstream section. |
| | 02 Aug | 1 | Drift sampling (T1) | Drift nets were collected after 24 hours, and macroinvertebrate samples were preserved with 96% ethanol for laboratory sorting. |
| | 05 Aug | 4 | Physical parameters | Conductivity, dissolved oxygen, pH, and temperature were measured in both upstream and downstream sections of each flume (background data, Figure S1). |
| | 06 Aug | 5 | Hydraulic parameters + light measurement | Flow velocity and water depth were measured in both upstream and downstream sections of each flume (background data, see Supplementary). Light intensity under the water surface (in lux) was checked and calibrated when needed (background data, Figure S1). |
| | 08 Aug | 7 | Drift nets placed | Drift nets were placed at the outlet of each upstream section. |
| | 09 Aug | 8 | Drift sampling (T2) | Drift nets were collected after 24 hours, and macroinvertebrate samples were preserved with 96% ethanol for laboratory sorting. |
| | 12 Aug | 11 | Physical parameters | Conductivity, dissolved oxygen, pH, and temperature were measured in both upstream and downstream sections of each flume (background data, Figure S1). |
| | 13 Aug | 12 | Hydraulic parameters + light measurement | Flow velocity and water depth were measured in both upstream and downstream sections of each flume (background data, Figure S1). Light intensity under the water surface (in lux) was checked and calibrated when needed (background data, Figure S1). |
| | 15 Aug | 14 | Drift nets placed | Drift nets were placed at the outlet of each upstream section. |
| | 16 Aug | 15 | Drift sampling (T3) | Drift nets were collected after 24 hours, and macroinvertebrate samples were preserved with 96% ethanol for laboratory sorting. |
| | 17 Aug | 16 | Physical parameters | Conductivity, dissolved oxygen, pH, and temperature were measured in both upstream and downstream sections of each flume (background data, Figure S1). |
| | 21 Aug | 20 | Hydraulic parameters + drift nets placed | Flow velocity and water depth were measured in both upstream and downstream sections of each flume (background data, Figure S1). Drift nets were placed at the outlet of each upstream section. |
| | 22 Aug | 21 | Drift sampling (T4) | Drift nets were collected after 24 hours, and macroinvertebrate samples were preserved with 96% ethanol for laboratory sorting. |
| | 23 Aug | 22 | End of the treatment phase + sampling of benthic macroinvertebrates + physical parameters | Benthic macroinvertebrate samples were collected in the designed spot (see Figure 2) in each section per flume. Preserved with 96% ethanol for laboratory sorting. Conductivity, dissolved oxygen, pH, and temperature were measured in both upstream and downstream sections of each flume (background data, Figure S1). |

**Table S2**. Common taxa* (abundance > 0.5%) used in generalized linear latent variable models (GLLVMs). Dispersal ability was assigned by combining different traits of the animals: drift propensity (Sarremejane et al., 2020), dispersal strategy, and locomotion (Tachet et al., 2010). For each taxon, we calculated active and passive disperser scores, and dispersal ability was assigned based on the highest score. All traits are provided with standardized fuzzy coding, with scoring based on the affinity of each taxon to the different modalities of the traits. Affinity scores sum up to one within each trait. Active and passive scores were calculated by averaging selected trait modalities, with the active score being the average of frequent drifter, aquatic active, aerial active, flier, surface swimmer, water swimmer, and crawler, whereas the passive score was the average of rare drifter, aquatic passive, aerial passive, burrower, interstitial, temporarily attached and permanently attached.

| **Order** | **Taxa** | **Drift propensity** | | | **Dispersal strategy** | | | | **Locomotion** | | | | | | | | **Active score** | **Passive score** | **Assigned dispersal ability** |
|---|---|---|---|---|---|---|---|---|---|---|---|---|---|---|---|---|---|---|---|
| | | **Rare** | **Occasional** | **Frequent** | **Aquatic passive** | **Aquatic active** | **Aerial passive** | **Aerial active** | **Flier** | **Surface swimmer** | **Water swimmer** | **Crawler** | **Burrower** | **Interstitial** | **Temporarily attached** | **Permanently attached** | | | |
| Amphipoda | *Gammarus sp.* | 0.17 | 0.50 | 0.33 | 0.60 | 0.40 | 0.00 | 0.00 | 0.00 | 0.00 | 0.33 | 0.50 | 0.00 | 0.17 | 0.00 | 0.00 | 0.22 | 0.13 | ACTIVE |
| Coleoptera | *Elmis sp.* | 0.60 | 0.40 | 0.00 | 0.40 | 0.20 | 0.00 | 0.40 | 0.17 | 0.00 | 0.00 | 0.67 | 0.00 | 0.17 | 0.00 | 0.00 | 0.20 | 0.17 | ACTIVE |
| Coleoptera | *Oreodytes sp.* | 0.75 | 0.25 | 0.00 | 0.33 | 0.17 | 0.00 | 0.50 | 0.14 | 0.00 | 0.43 | 0.43 | 0.00 | 0.00 | 0.00 | 0.00 | 0.24 | 0.15 | ACTIVE |
| Diptera | Ceratopogonidae | 1.00 | 0.00 | 0.00 | 0.50 | 0.17 | 0.00 | 0.33 | 0.00 | 0.13 | 0.38 | 0.13 | 0.38 | 0.00 | 0.00 | 0.00 | 0.16 | 0.27 | PASSIVE |
| Diptera | Chironomini | 0.33 | 0.50 | 0.17 | 0.29 | 0.14 | 0.43 | 0.14 | 0.00 | 0.00 | 0.11 | 0.33 | 0.22 | 0.11 | 0.22 | 0.00 | 0.13 | 0.23 | PASSIVE |
| Diptera | Orthocladiinae | 0.33 | 0.50 | 0.17 | 0.40 | 0.20 | 0.20 | 0.20 | 0.00 | 0.00 | 0.13 | 0.38 | 0.13 | 0.25 | 0.13 | 0.00 | 0.15 | 0.20 | PASSIVE |
| Diptera | Simuliidae | 0.00 | 0.40 | 0.60 | 0.25 | 0.25 | 0.38 | 0.13 | 0.00 | 0.00 | 0.00 | 0.29 | 0.00 | 0.14 | 0.57 | 0.00 | 0.18 | 0.19 | PASSIVE |
| Diptera | Tanypodinae | 0.33 | 0.50 | 0.17 | 0.40 | 0.20 | 0.20 | 0.20 | 0.00 | 0.00 | 0.43 | 0.29 | 0.14 | 0.14 | 0.00 | 0.00 | 0.18 | 0.17 | ACTIVE |
| Diptera | Tanytarsini | 0.33 | 0.50 | 0.17 | 0.17 | 0.17 | 0.50 | 0.17 | 0.00 | 0.00 | 0.25 | 0.38 | 0.00 | 0.13 | 0.25 | 0.00 | 0.16 | 0.20 | PASSIVE |
| Ephemeroptera | *Baetis sp.* | 0.00 | 0.25 | 0.75 | 0.33 | 0.22 | 0.11 | 0.33 | 0.00 | 0.00 | 0.38 | 0.50 | 0.00 | 0.13 | 0.00 | 0.00 | 0.31 | 0.08 | ACTIVE |
| Ephemeroptera | *Ephemerella sp.* | 0.33 | 0.50 | 0.17 | 0.22 | 0.33 | 0.11 | 0.33 | 0.00 | 0.00 | 0.17 | 0.83 | 0.00 | 0.00 | 0.00 | 0.00 | 0.26 | 0.10 | ACTIVE |
| Plecoptera | *Leuctra sp.* | 0.33 | 0.50 | 0.17 | 0.40 | 0.40 | 0.00 | 0.20 | 0.00 | 0.00 | 0.00 | 0.63 | 0.25 | 0.13 | 0.00 | 0.00 | 0.20 | 0.16 | ACTIVE |
| Plecoptera | *Protonemura sp.* | 0.25 | 0.50 | 0.25 | 0.40 | 0.40 | 0.00 | 0.20 | 0.00 | 0.00 | 0.00 | 1.00 | 0.00 | 0.00 | 0.00 | 0.00 | 0.26 | 0.09 | ACTIVE |
| Sphaeriida | *Pisidium sp.* | 1.00 | 0.00 | 0.00 | 0.50 | 0.25 | 0.25 | 0.00 | 0.00 | 0.00 | 0.00 | 0.13 | 0.50 | 0.13 | 0.25 | 0.00 | 0.05 | 0.38 | PASSIVE |
| Tricladida | *Polycelis sp.* | 1.00 | 0.00 | 0.00 | 0.33 | 0.67 | 0.00 | 0.00 | 0.00 | 0.14 | 0.00 | 0.71 | 0.00 | 0.14 | 0.00 | 0.00 | 0.22 | 0.21 | ACTIVE |
| Trombidiformes | Hydrachnidia | Traits data not available. Dispersal ability assigned based on (Di Sabatino et al., 2000) and (Smith et al., 2010) | | | | | | | | | | | | | | | 0.00 | 0.00 | PASSIVE |
| Tubificida | Naididae | 0.75 | 0.27 | 0.04 | 0.79 | 0.21 | 0.00 | 0.00 | 0.00 | 0.00 | 0.33 | 0.00 | 0.26 | 0.31 | 0.07 | 0.02 | 0.08 | 0.31 | PASSIVE |

* *Note*: relative abundance was calculated within each dataset (drift upstream, benthos upstream, drift downstream, and benthos downstream).

**Table S3**. Full taxa list with abundances for drifting and benthic macroinvertebrates. Organisms exclusive of either drift or benthos are marked in bold.

| GROUP | TAXA | TOTAL | DRIFT | BENTHOS |
|---|---|---|---|---|
| Coleoptera | *Elmis sp.* | 106 | 81 | 25 |
| Coleoptera | *Elodes sp.* | 22 | 21 | 1 |
| **Coleoptera** | ***Esolus sp.*** | **1** | **0** | **1** |
| Coleoptera | *Hydraena sp.* | 6 | 5 | 1 |
| **Coleoptera** | ***Hydroporus sp.*** | **2** | **2** | **0** |
| Coleoptera | *Laccophilus sp.* | 6 | 4 | 2 |
| Coleoptera | *Limnius sp.* | 53 | 14 | 39 |
| Coleoptera | *Oreodytes sp.* | 52 | 14 | 38 |
| **Crustacea** | ***Asellus sp.*** | **1** | **0** | **1** |
| Crustacea | *Gammarus sp.* | 2600 | 679 | 1921 |
| Diptera | Ceratopogonidae | 54 | 4 | 50 |
| Diptera | Chironomini | 592 | 24 | 568 |
| **Diptera** | **Culicidae** | **1** | **0** | **1** |
| **Diptera** | **Dixidae** | **4** | **4** | **0** |
| Diptera | Empididae | 52 | 5 | 47 |
| Diptera | Limoniidae | 33 | 3 | 30 |
| Diptera | Orthocladiinae | 6253 | 1638 | 4615 |
| Diptera | Psychodidae | 3 | 1 | 2 |
| **Diptera** | **Ptychopteridae** | **8** | **0** | **8** |
| Diptera | Simuliidae | 1499 | 1164 | 335 |
| Diptera | Stratiomyidae | 4 | 3 | 1 |
| **Diptera** | **Tabanidae** | **16** | **0** | **16** |
| Diptera | Tanypodinae | 689 | 42 | 647 |
| Diptera | Tanytarsini | 1499 | 150 | 1349 |
| Ephemeroptera | *Baetis sp.* | 1092 | 517 | 575 |
| Ephemeroptera | *Ecdyonurus sp.* | 15 | 6 | 9 |
| Ephemeroptera | *Ephemerella sp.* | 425 | 383 | 42 |
| Ephemeroptera | Leptophlebiidae | 8 | 5 | 3 |
| **Megaloptera** | ***Sialis*** *sp.* | **3** | **0** | **3** |
| **Mollusca** | ***Pisidium*** *sp.* | **61** | **0** | **61** |
| Nematoda | Mermithidae | 6 | 1 | 5 |
| **Odonata** | ***Cordulegaster*** *sp.* | **2** | **0** | **2** |
| **Oligochaeta** | **Lumbricidae** | **2** | **2** | **0** |
| Oligochaeta | Naididae | 43 | 26 | 17 |
| **Plecoptera** | ***Chloroperla*** *sp.* | **1** | **0** | **1** |
| Plecoptera | *Leuctra sp.* | 284 | 110 | 174 |
| Plecoptera | Nemouridae | 932 | 283 | 649 |
| **Plecoptera** | ***Perlodes*** *sp.* | **3** | **3** | **0** |
| **Plecoptera** | ***Protonemura*** *sp.* | **192** | **192** | **0** |
| **Trichoptera** | **Brachicentridae** | **2** | **2** | **0** |
| **Trichoptera** | ***Hydropsyche*** *sp.* | **4** | **4** | **0** |
| Trichoptera | Limnephilidae | 28 | 21 | 7 |
| **Trichoptera** | ***Lithax*** *sp.* | **1** | **0** | **1** |
| **Trichoptera** | ***Odontocerum*** *sp.* | **1** | **1** | **0** |
| **Trichoptera** | ***Philopotamus*** *sp.* | **4** | **4** | **0** |
| Trichoptera | *Rhyacophila sp.* | 12 | 10 | 2 |
| Trichoptera | *Sericostoma sp.* | 28 | 1 | 27 |
| Tricladida | *Polycelis sp.* | 52 | 33 | 19 |
| Trombidiformes | Hydrachnidia | 133 | 102 | 31 |

**Table S4.** Roadmap of the data analysis, including their matching aim and the target variable/community. The structure of the table matches how analyses are reported in the data analysis section of the manuscript.

| Aim | Target variables | Target dataset | Model | Fixed factors | Random factors |
|---|---|---|---|---|---|
| Aim 1: local effects of stressors | Metrics (taxa richness and abundance) | Drift | Linear mixed models (from *lme4* R package) | Drought (categorical; 0, 1), light (categorical; 0, 1) and their interaction, plus time (categorical; T1, T2, T3, and T4) | Flume nested within header tank |
| | | Benthos | | Drought (categorical; 0, 1), light (categorical; 0, 1), and their interaction | |
| Aim 2: downstream effect of stressors | | Drift | | Stressors arrangement (categorical; control, separated, and combined) plus time | |
| | | Benthos | | Stressors arrangement (categorical; control, separated, and combined) | |
| Aim 1: local effects of stressors | Community structure (common taxa) | Drift | Generalized linear latent variable models (from *gllvm* R package) | Drought (categorical; 0, 1), light (categorical; 0, 1) and their interaction, plus time (categorical; T1, T2, T3, and T4) | Flume nested within header tank |
| | | Benthos | | Drought (categorical; 0, 1), light (categorical; 0, 1), and their interaction | |
| Aim 2: downstream effect of stressors | | Drift | | Stressors arrangement (categorical; control, separated, and combined) plus time | |
| | | Benthos | | Stressors arrangement (categorical; control, separated, and combined) | |

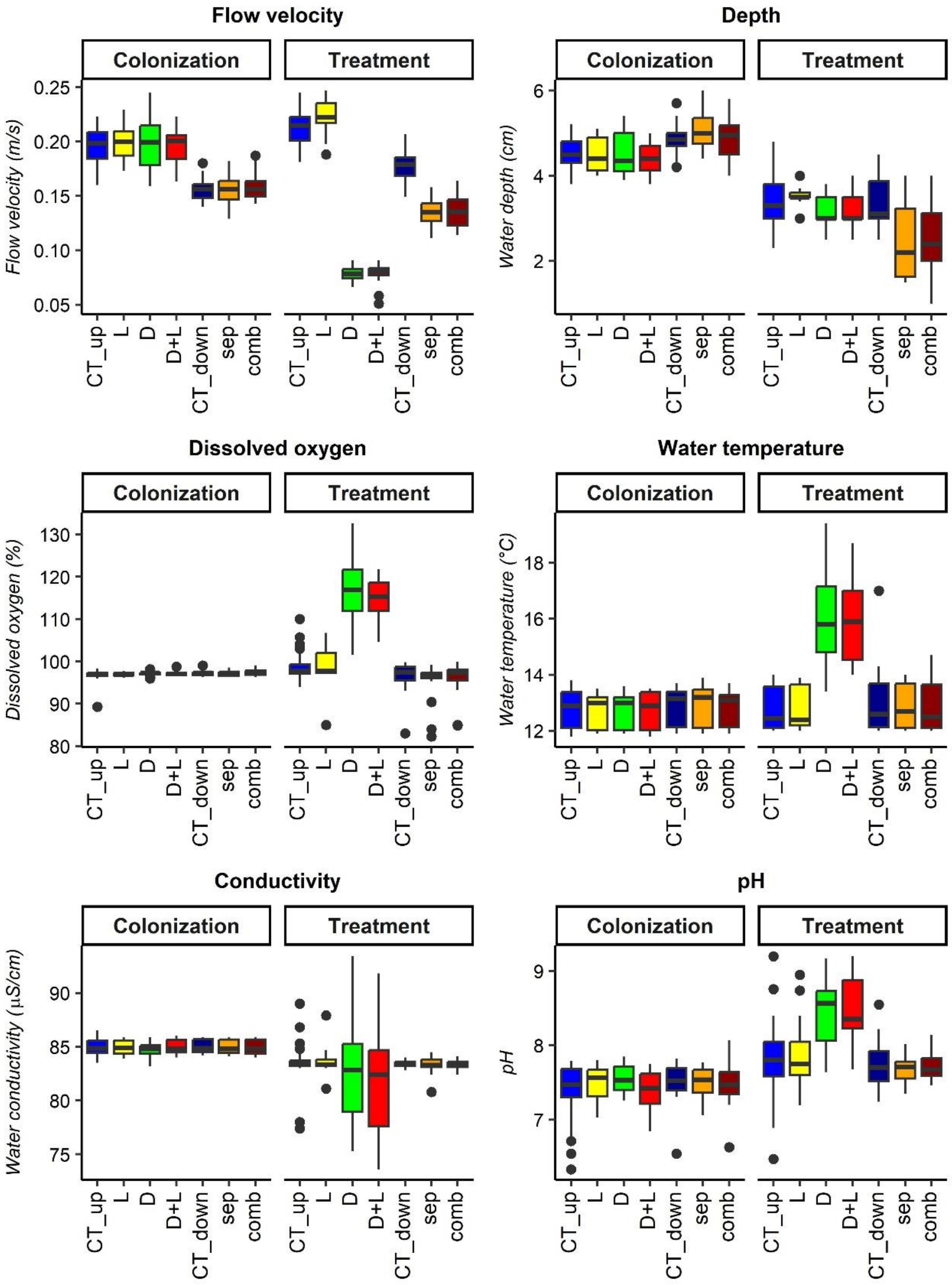


**Figure S1.** Boxplots summarizing the background data (flow meter, depth, and multiparametric probe) for each treatment level during both the colonization and treatment phases. Note that for each plot, the first four boxes refer to the upstream sections (CT_up = control upstream, L = light treatment, D = drought treatment, D+L = drought and light treatment) and the last three to the downstream ones (CT_down = control downstream, sep = separated treatment, comp = combined treatment).

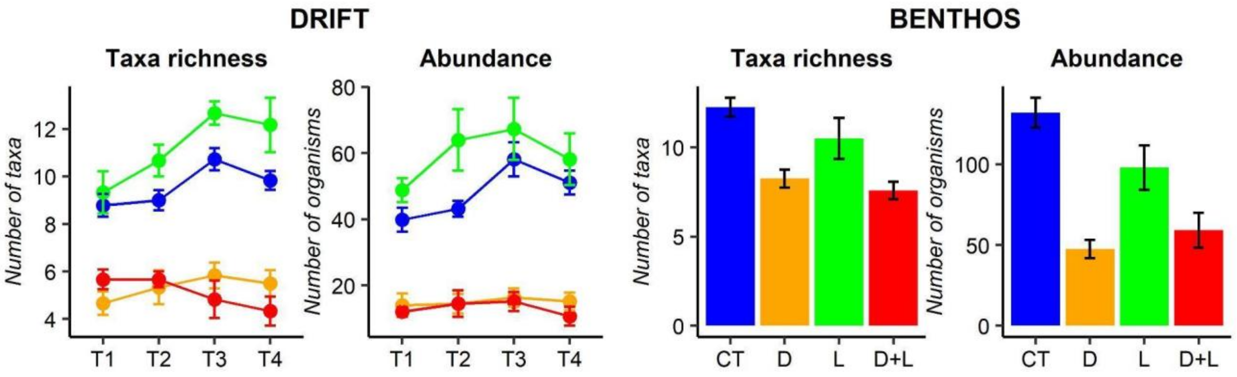


**Figure S2**. Variability of taxa richness and abundance metrics for drifting and benthic macroinvertebrates in upstream sections. T1-4 refers to the weekly sampling of drifting macroinvertebrates. CT = control, D = drought treatment, L = light treatment, D+L = drought and light treatment.

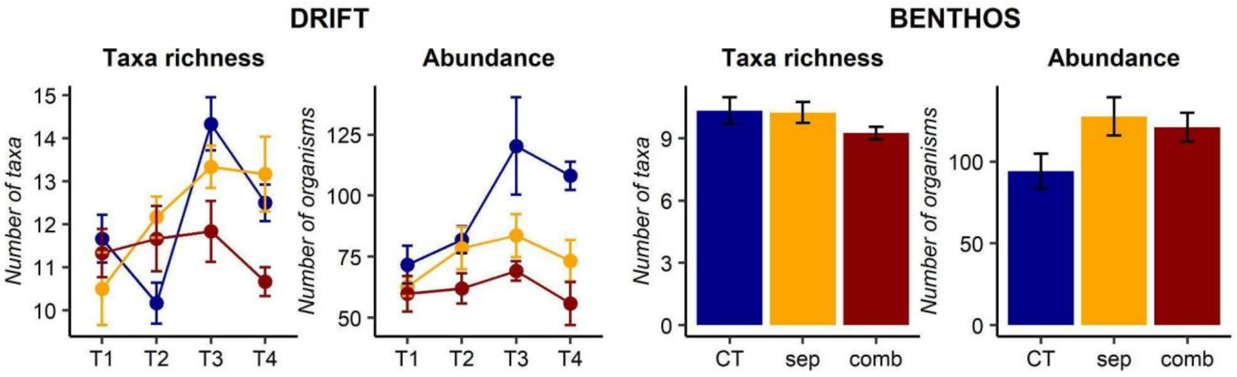


**Figure S3.** Variability of taxa richness and abundance metrics for drifting and benthic macroinvertebrates in downstream sections. T1-4 refers to the weekly sampling of drifting macroinvertebrates. CT = control (downstream), sep = stressors separated in upstream tributaries, comb = stressors combined in upstream tributaries.
